# stFormer: a foundation model for spatial transcriptomics

**DOI:** 10.1101/2024.09.27.615337

**Authors:** Shenghao Cao, Kaiyuan Yang, Jiabei Cheng, Jiachen Li, Hong-Bin Shen, Xiaoyong Pan, Ye Yuan

## Abstract

Recent foundation models for single-cell transcriptomics data generate informative, context-aware representations for genes and cells. The **S**patial **T**ranscriptomics (ST) data offer extra positional insights, which were not considered by these single-cell models. Here, we introduce stFormer, a transformer model tailored for ST data. stFormer employs the cross-attention module to incorporate spatial ligand genes into the transformer encoder of single-cell transcriptomics. To unify different ST technologies with trade-off between resolution and gene coverage, we propose a biased cross-attention method that enables the model to do learning with single-cell resolution on whole-transcriptome but low-resolution Visium data. We collected human Visium datasets from a public ST database and performed cell type deconvolution, generating ~4.1 million pretraining samples. As a foundation model for ST data, stFormer improved performance upon the state-of-the-art single-cell foundation model, scFoundation, across a variety of tasks, including cell clustering, batch effect correction, cell type prediction, and gene function prediction. stFormer also revealed intercellular ligand-receptor signaling responses via *in silico* perturbation.

## Introduction

The large language model based on the **G**enerative **P**retrained **T**ransformer (GPT) architecture has made groundbreaking progress in the field of artificial intelligence^1,2^. Recently, the transformer pretraining method was applied to building foundation models for single-cell transcriptomics data^3-5^. The foundation models generated contextual gene and cell embeddings for single cells. These models have demonstrated strong performance across a range of downstream applications. However, despite their power in capturing complex gene and cell features, they are fundamentally constrained to modeling intracellular transcriptomic signals.

Notably, the intracellular transcription is not only regulated by intracellular gene networks, but also mediated by extracellular microenvironment, which is absent from single-cell transcriptomics data. ST technologies address this limitation by profiling gene expressions in their native tissue locations. Previous transformer models for single-cell data have not taken advantage of these spatial information.

To bridge this gap, we developed stFormer, a transformer model adapted for ST data to integrate both intracellular and spatial contexts. The architecture design of stFormer presents two key innovations. First, stFormer introduces the cross-attention mechanism, previously unexplored in single-cell models, for capturing ligand-mediated gene interactions. Second, stFormer proposes a biased cross-attention method which enables single-cell resolution learning from cell-type deconvolved visium data, a widely available spatial resource with whole-transcriptome gene coverage.

We assembled a pretraining corpus comprising ~4.1 million spatial samples from public human Visium datasets, spanning diverse tissues, development stages, and disease states. As a foundation model for ST data, stFormer demonstrated strong generalization capabilities. It generated spatially aware embeddings that outperformed those from single-cell models in both cell-level and gene-level prediction tasks. Furthermore, stFormer enables *in silico* perturbation of spatial ligand–receptor interactions, expanding its utility for mechanistic exploration.

## Results

### stFormer architecture and pretraining

stFormer harnesses the transformer architecture to capture gene interactions for optimizing predictive accuracy of self-supervised masked learning objective. The key improvement over the previous single-cell models is that we insert a cross-attention module after the self-attention module in the transformer block to compute attentions with ligand genes from niche cells (**Fig. 1a and Methods**). In this way, stFormer integrates cell-cell communication effects via ligand secretion and reception into intracellular gene embeddings.

**Fig. 1.**
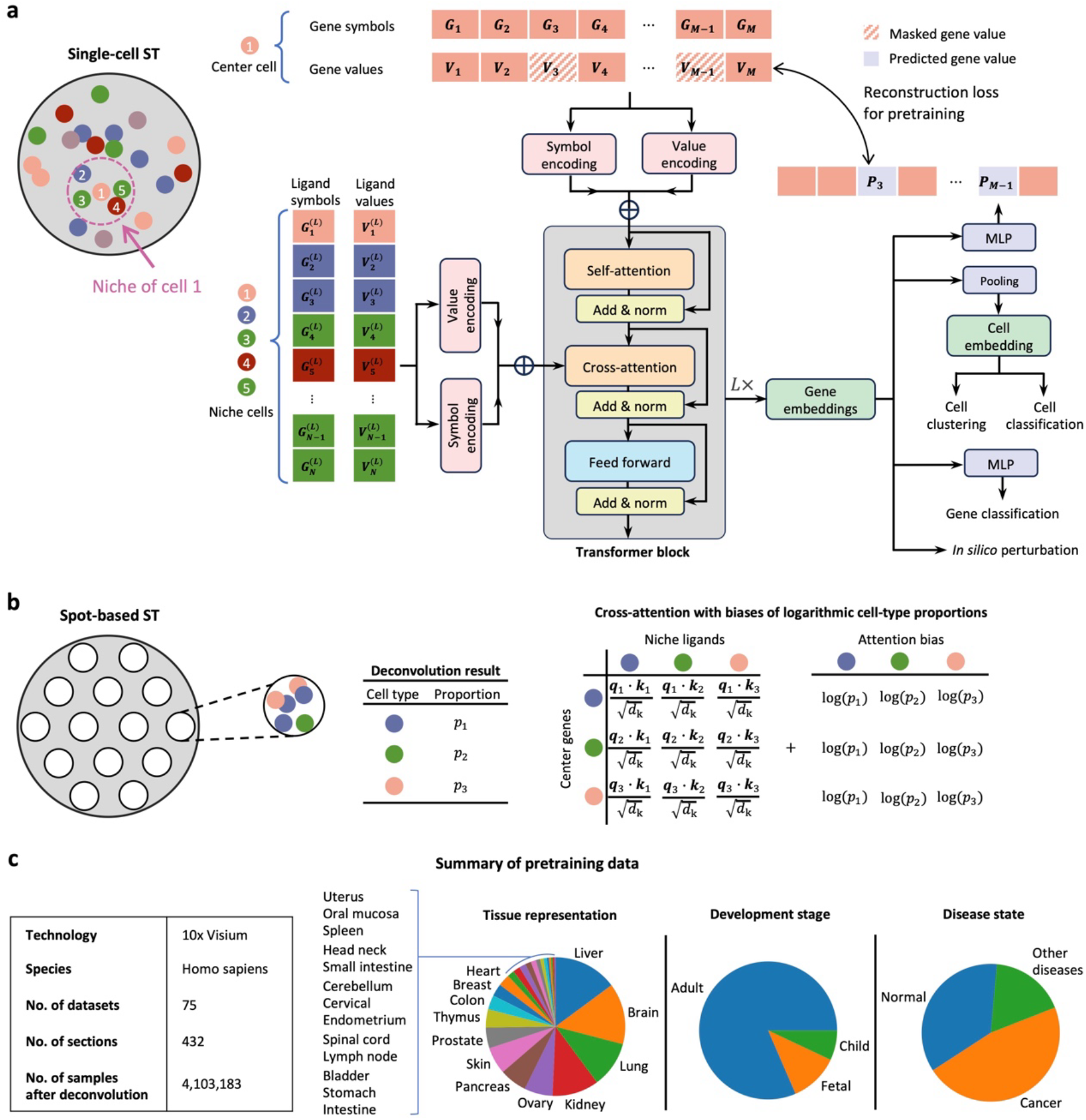
Model schematic. **a**, For the spatial transcriptomics (ST) data with single-cell resolution, we regard each cell as a centric node and the neighborhood within a preset radius as its niche. An input sample comprises gene symbols and values of the center cell as well as ligand gene symbols and values of the niche cells. The samples are fed into a stack of *L* = 6 identical transformer block layers and output the embeddings of genes in the center cell. The model parameters are pretrained by minimizing the reconstruction loss of the masked gene values. Then, the gene embeddings can serve as input for multiple downstream tasks. **b**, For the ST data based on spot RNA sequencing, we deconvolve the cell-type composition at each spot first. Then, adding a bias of logarithmic cell-type proportion to each attention score in cross-attention can unify the case within the single-cell framework in **a**. See **Methods** for details. **c**, Summary of pretraining data, totally about 4.1 million samples at cell-type resolution. Pie charts display the proportional data distribution, categorized by tissue origin, development stage, and disease state, respectively.

A variety of ST technologies are available with a trade-off between resolution and gene coverage. One of the most commonly used platforms is the 10x Genomics Visium^6^. It captures whole transcriptomes at the arranged spots in the tissue section. However, the spot diameter, i.e. the resolution, is 55 *μ*m, which is several-fold larger than a typical cell size, so each spot may contain multiple cells. Deconvolution methods such as cell2location^7^ can estimate the cell-type proportions and cell-type specific expressions at each spot, but the cell number and cell specific expressions remain unknown. To address this problem, we propose the cell-type-wise biased cross-attention method, which enables the model to do single-cell resolution learning on spot-resolution Visium data (**Fig. 1b**). The theoretical details and derivation are provided in the **Methods** section.

We collected 75 human Visium datasets comprised of 432 tissue sections from CROST^8^, a comprehensive repository of ST data, and complemented them with paired single-cell transcriptomics data (see details in **Supplementary Data 1**). Then we deconvolved these Visium datasets with their paired single-cell reference, generating ~4.1 million spatial samples at cell-type resolution. These samples represented heterogeneous tissues, development stages, and disease states (**Fig. 1c**). Finally, we pretrained stFormer on these samples through self-supervised learning, which predicted masked gene expressions based on visible genes and spatial ligand genes (**Methods**).

### stFormer improved batch removal and cell clustering performance

For model evaluation, 0.5% of pretraining samples were randomly held out as test data. Both training and test loss for the masked learning objective followed power-law decays with increasing sample size (**Fig. 2a and Supplementary Figure S1**). After pretraining, we evaluated the batch effect on cell embeddings produced by stFormer using a public benchmarking dataset^9,10^, a single-cell RNA-seq project including 8 patients and 3 control subjects. The single-cell transcriptomics data were imported into stFormer at the setting of single-cell mode, where intercellular cross-attention module was disabled (**Methods**). UMAP visualizations revealed that compared to scFoundation^5^, stFormer-derived cell embeddings improved batch mixing, as quantified by a higher ASW_batch_ score (a batch correction metric^11^), while maintained comparable cell type separation (**Fig. 2b**). After fine-tuning, stFormer showed a modest decline in batch correction metric, whereas its cell type separation improved markedly (**Supplementary Figure S2**).

**Fig. 2.**
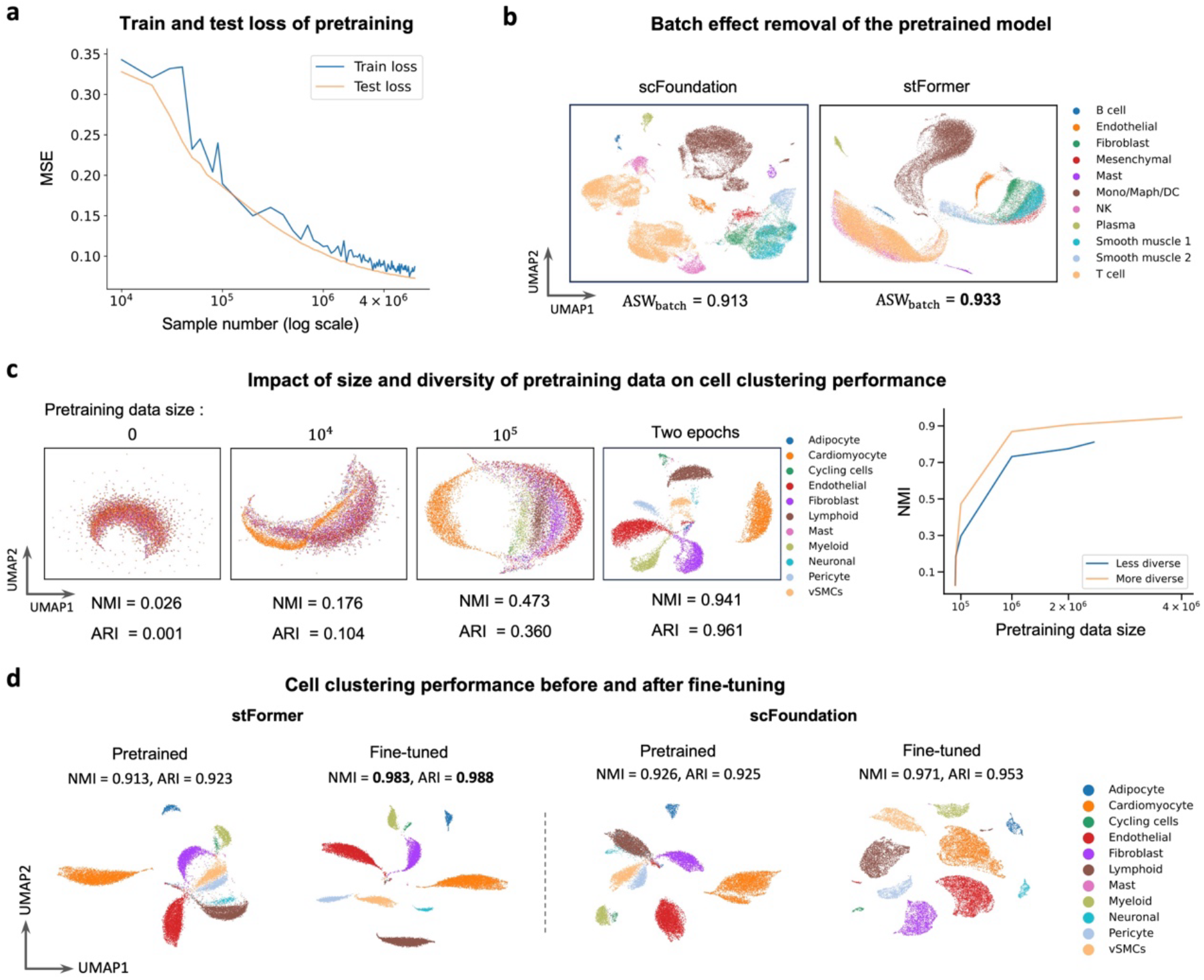
Cell clustering performance. **a**, Train and test loss of the pretraining objective as the number of pretraining samples increased. MSE, mean squared error. **b**, UMAP visualizations of cell embeddings generated by scFoundation and stFormer for an scRNA-seq benchmarking dataset, with points colored by cell types. stFormer removed batch effects in the data while preserving biological variance. Batch correction metric ASW_batch_ is indicated below each UMAP plot. **c**, Clustering performance on a test Visium data using cell embeddings generated from stFormer models pretrained on different data sizes and diversities. The left four plots display UMAP visualizations of the pretrained cell embeddings, with points colored by cell types. The pretraining data sizes are indicated above the UMAP plots, and two clustering evaluation metrics (NMI and ARI) are shown below. The right plot displays the NMI variation with pretraining data size at two diversity levels. **d**, Clustering performance of cell embeddings generated from pretrained and fine-tuned stFormer on a test Visium data, compared with scFoundation.

We further evaluated stFormer’s cell clustering performance using its cell embeddings on a myocardial Visium section^12^, which was unseen in the pretraining data. We found that as pretraining data scaled up, UMAP visualizations^13^ of stFormer-derived cell embeddings showed progressively clearer separation, and both NMI (normalized mutual information) and ARI (adjusted Rand index) demonstrated improved alignment with cell type annotations (**Fig. 2c**). At the end of pretraining (two epochs), the resulting clusters closely matched ground-truth annotations, as evidenced by high NMI (0.941) and ARI (0.961) scores. Furthermore, we pretrained an alternative stFormer instance on a subset of Visium datasets (55 out of 75), yielding ~2.4 million spatial samples. The subset data represented a reduced biological diversity. As a result, although NMI also increased with sample size, it consistently underperformed stFormer-4.1M and exhibited inferior asymptotic performance (**Fig. 2c and Supplementary Figure S3**).

We evaluated the cell clustering performance after fine-tuning. Specifically, we fine-tuned the pretrained model using the self-supervised masked learning objective on a test Visium section. In this way, all parameters were adapted to this test data. Then we clustered the cell embeddings produced by the fine-tuned model on the test data. As shown in **Fig. 2d**, NMI and ARI metrics for stFormer and scFoundation both improved after fine-tuning. Although stFormer initially underperformed relative to scFoundation, likely due to its more limited pretraining data size (~4.1 million vs ~50 million), it ultimately surpassed scFoundation following fine-tuning.

### Spatially aware cell embeddings improved cell type predictions

We appended a three-layer perceptron classifier to stFormer for cell type prediction based on cell embeddings (**Methods**). Since cell type annotations were already available through deconvolution for Visium data, there was no need to predict cell types for them. Therefore, we tested on a single-cell ST data, a CosMx SMI (**S**patial **M**olecular **I**mager) dataset from human pancreas tissue^14^. This dataset covered over 18,000 genes using a whole-transcriptome panel. stFormer offers an option— *max number of niche cells*—to define the radius of spatial niche where ligand genes are taken as input (**Methods**).

To eliminate class imbalance effects, we equalized the sample sizes by downsampling each cell type. Then we evaluated stFormer via fivefold cross-validation under distinct niche radius settings, and compared with scFoundation^5^. The last two transformer layers along with the classifier were fine-tuned on the training stages, while all other layers were frozen. We evaluated two stFormer versions. One is stFormer-4.1M, which was pretrained on the entire corpus comprising ~4.1 million samples; the other is stFormer-2.4M, which was pretrained on a corpus subset compring ~2.4 million samples.

As can be seen in **Fig. 3a**, the predictive accuracy of both stFormer versions generally followed a trend of first increasing and then stabilizing as niche radius expanded. stFormer-4.1M consistently outperformed stFormer-2.4M and scFoundation at all settings of niche radius, while stFormer-2.4M initially underperformed scFoundation and then surpassed it as *max number of niche cells* exceeded 15. When *max number of niche cells* was set to 20, stFormer-4.1M achieved nearly optimal median accuracy with relatively low variance. At this point, stFormer-4.1M demonstrated superior prediction accuracy across a greater number of cell types compared to scFoundation as shown in **Fig. 3b**.

**Fig. 3.**
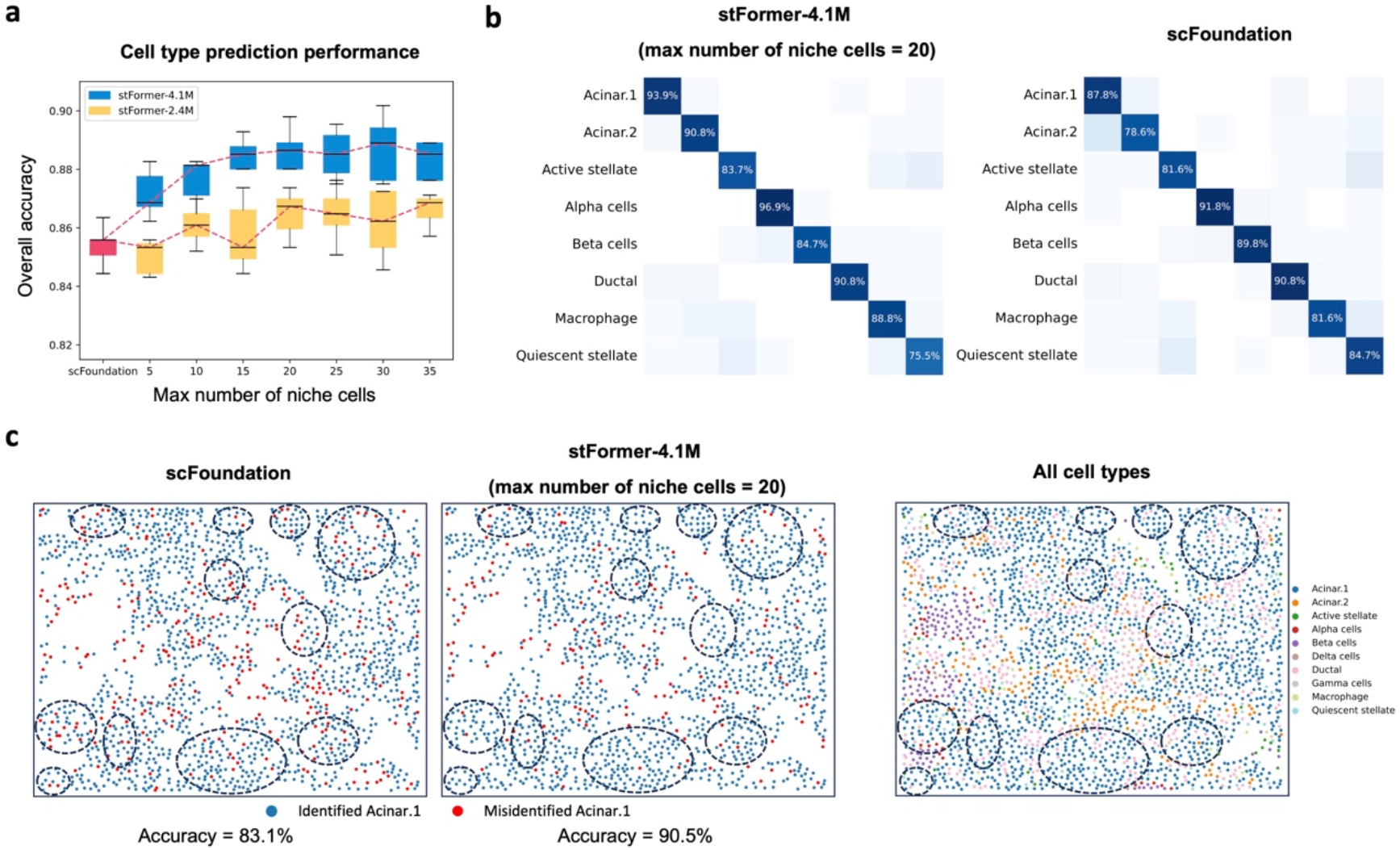
Cell type prediction performance fine-tuned on a pancreas single-cell ST data. **a**, Predictive accuracy by fivefold cross-validation of two stFormer versions at different settings of *max number of niche cells*, compared to scFoundation. *Max number of niche cells* is an option in stFormer determining the spatial niche radius. stFormer-4.1M was pretrained on ~4.1 million samples while stFormer-2.4M was pretrained on ~2.4 million samples. **b**, Median confusion matrices by fivefold cross-validation comparing the predictive results from stFormer-4.1M (*max number of niche cells* = 20) and scFoundation. **c**, The left two plots compare the spatial distributions of correctly and incorrectly classified acinar.1 cells between scFoundation and stFormer on an excluded FOV. Regions with improved predictions are circled. The right plot displays cell type distributions in the circled regions, which exhibit similar spatial patterns.

To visualize the spatial distribution of predictions, we excluded one field-of-view (FOV) from the dataset, and downsampled remaining data to balanced class distribution. Then, we split training (80%) and validation (20%) sets, and performed model training and selection on them respectively. Finally, we predicted cell types on the held-out FOV. stFormer-4.1M (*max number of niche cells* = 20) outperformed scFoundation with an overall accuracy of 88.6% compared to 84.4% (**Supplementary Figure S4**). Acinar.1 cells accounted for the largest proportion (63.6%) of cells in the FOV. We compared the spatial distributions of acinar.1 cell predictions from scFoundation and stFormer-4.1M in **Fig. 3c**. The circled regions highlighted acinar.1 cells with improved predictions by stFormer. These areas exhibited similar spatial patterns of cell type distributions.

### Spatially aware gene embeddings improved gene function predictions

We appended a three-layer perceptron classifier to stFormer for gene function prediction based on gene embeddings (**Methods**). We first took the TGF-beta signaling pathway as an example. The TGF-beta signaling pathway is initiated by the binding of TGF-beta ligands to their receptors. We adopted the KEGG gene set “TGF-beta signaling pathway” provided by MSigDB^15^, where external signaling genes were excluded. An equal number of genes outside this set were randomly sampled as negative controls. It has been reported that the TGF-beta signaling pathway showed increased activity in fibrotic myocardium^12^, so we evaluated stFormer on a Visium section of fibrotic myocardium^12^ via fivefold cross-validation, and compared with scFoundation. The last two transformer layers and the classifier were fine-tuned on the training sets for predicting gene inclusion within the gene set, with performance assessed on the validation sets. As shown in **Fig. 4a and b**, the median ROC (receiver operating characteristic) curve and PR (precision recall) curve of stFormer both outperformed those of scFoundation, with median AUROC (area under the ROC curve) of 0.90 compared to 0.60 and median AUPRC (area under the PR curve) of 0.85 compared to 0.65.

**Fig. 4.**
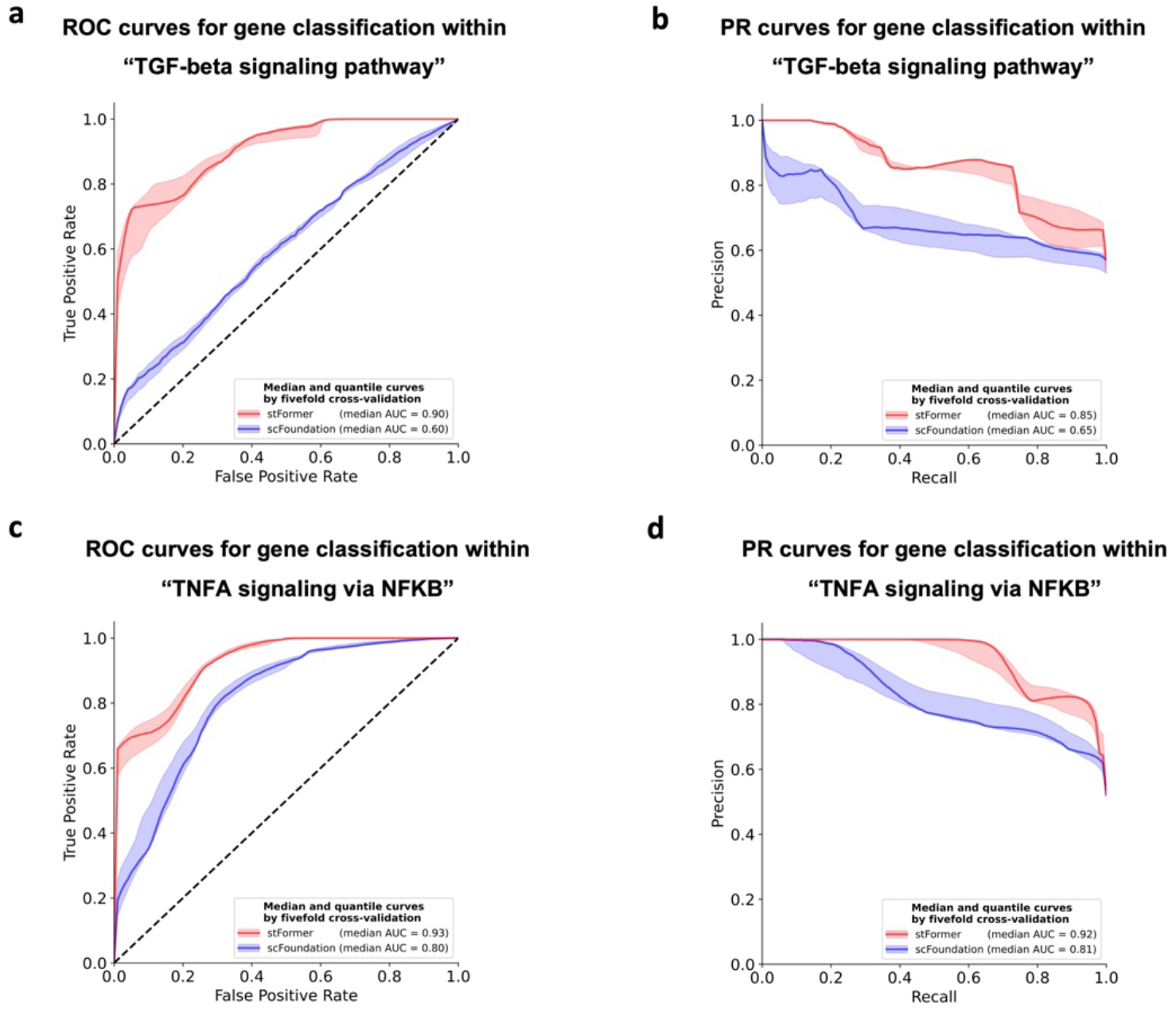
Gene function prediction performance fine-tuned on a myocardium Visium data. **a**,**b**,**c**,**d**, ROC curves (**a**) and PR curves (**b**) by fivefold cross-validation for predicting genes within gene set “TGF-beta signaling pathway”. ROC curves (**c**) and PR curves (**d**) by fivefold cross-validation for predicting genes within gene set “TNFA signaling via NFKB”. The evaluations were performed on a fibrotic myocardial Visium section. Red and blue lines represent median curves for stFormer and scFoundation respectively, and shaded regions represent 40% to 60% quantile intervals.

As the NFκB pathway was also found to be enriched in myocardial fibrosis^12^, we additionally tested gene function prediction using the “TNFA signaling via NFKB” hallmark gene set provided by MSigDB. As shown in **Fig. 4c and d**, the median ROC (receiver operating characteristic) curve and PR (precision recall) curve of stFormer also outperformed those of scFoundation, with median AUROC (area under the ROC curve) of 0.93 compared to 0.80 and median AUPRC (area under the PR curve) of 0.92 compared to 0.81.

### *In silico* perturbation revealed ligand-receptor signaling responses

Finally, we tested whether stFormer-derived gene embeddings captured responses of cell-cell signaling via ligand-receptor interactions. It was reported that the WNT signaling pathway plays a pivotal role in cardiac remodeling^16^. Thus, we chose a Visium section from ischemic myocardium^12^ and *in silico* reduced the gene expressions of WNT ligands and receptors specifically in cardiomyocytes (**Methods**). The deleterious effects were predicted by alterations of gene embeddings. We evaluated WNT signaling responses by examining the deleterious effects on housekeeping genes and WNT target genes, which we use direct targets listed on the Stanford Wnt homepage (https://wnt.stanford.edu/target_genes). As shown in **Fig. 5**, *In silico* downregulation of WNT ligands and receptors, either individually or combinedly, had a significantly stronger effect on WNT targets than on randomly selected genes, whereas housekeeping genes remained largely unaffected (**Fig. 5a-c**). Furthermore, combined downregulation of WNT ligands and receptors demonstrated significantly enhanced effects on WNT targets compared to individual downregulation of either ligands or receptors (**Fig. 5d**).

**Fig. 5.**
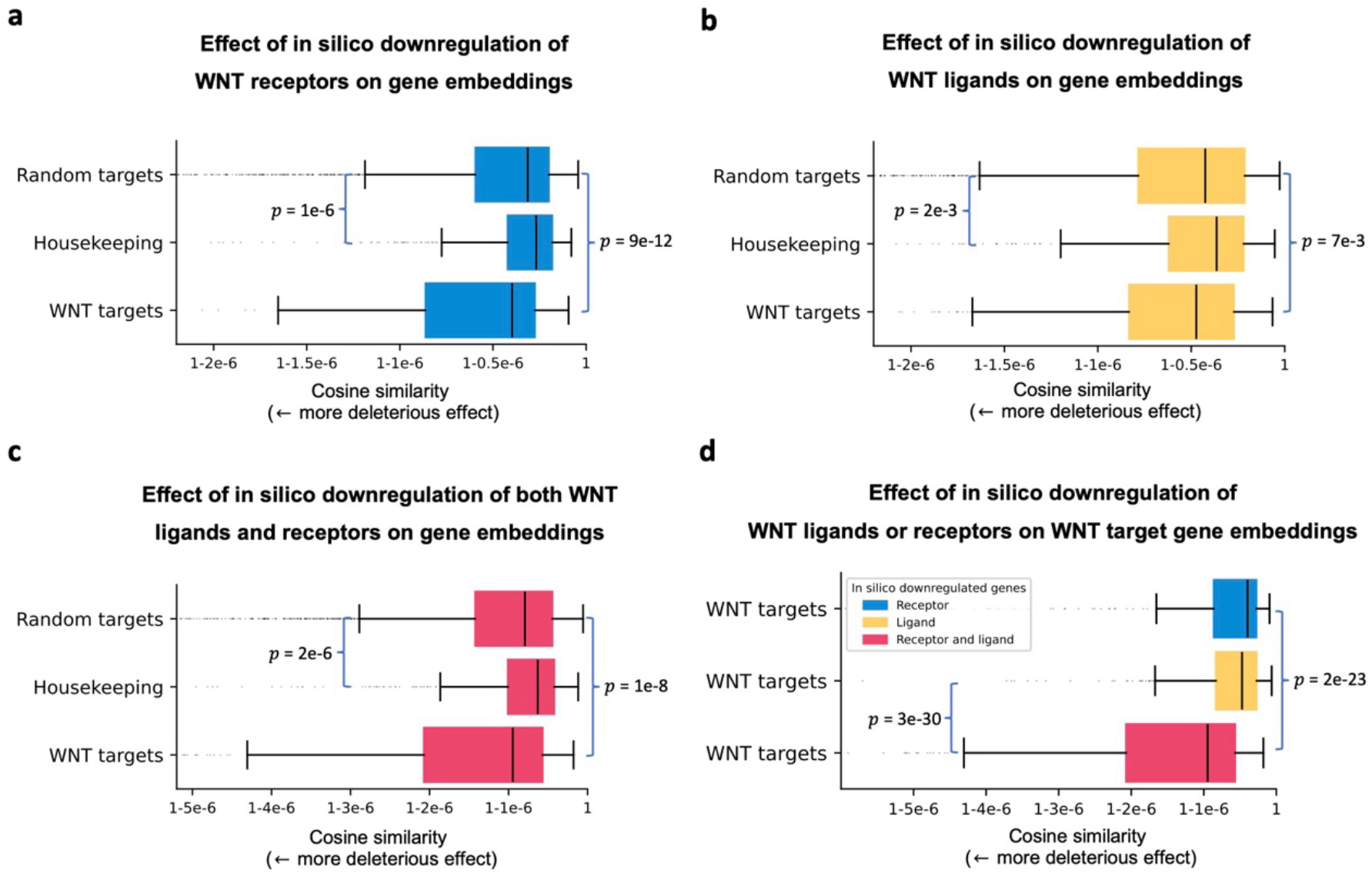
*In silico* perturbation revealed ligand-receptor signaling responses. **a**,**b**,**c** Effect of *in silico* downregulation of WNT receptors (**a**), WNT ligands (**b**), or both WNT ligands and receptors (**c**) on gene embeddings generated from pretrained stFormer. Target genes of the WNT signaling pathway were impacted significantly more than randomly selected genes, while housekeeping genes were significantly less impacted. (**d**) *In silico* downregulation of WNT ligands and receptors together impacted on WNT targets significantly more than separate downregulation of ligands or receptors. These tests were performed on an ischemic myocardium Visium section. FDR-corrected Wilcoxon p-values are shown beside.

## Discussion

Advances in large language models have inspired the development of foundation models for single-cell transcriptomics data^3-5^. The spatially resolved transcriptomics (ST) technologies, as newly emerging tools for transcriptome quantification, profile gene expressions while preserving spatial position information. However, the extra positional information is not considered in existing single-cell foundation models. Building foundation models for ST data presents two main challenges. One is how to incorporate spatial information into gene expression sequence model; the other is how to accommodate diverse ST technologies, which often involve trade-offs between resolution and gene coverage.

In this study, we introduce stFormer as a solution to the aforementioned challenges. stFormer harnesses the transformer architecture^1^. In addition to the self-attention module, which is widely adopted by the existing single-cell models to encode intracellular gene interactions, stFormer innovatively employs the cross-attention module to encode ligand-mediated intercellular gene interactions. In this way, stFormer incorporates ligand gene expressions in the spatially neighboring cells into single-cell transcriptomics. Furthermore, we proposed the biased cross-attention method which enables single-cell resolution learning from cell-type resolution visium data, a widely available spatial resource with whole-transcriptome gene coverage. We assembled ~4.1 million spatial samples from public human Visium datasets. After pretrained on these samples, stFormer is compatible with both single-cell and spot resolution ST data.

We demonstrated that after pretraining stFormer achieved generalization capabilities across diverse tasks. Pretraining via the masked gene prediction task highlighted stFormer’s core understanding of inter-gene associations. The cell clustering task revealed that pooling gene embeddings within single cells could generate cell embeddings that encode the characteristics of cell states. We further showed that stFormer removed batch effects in the cell embeddings through an scRNA-seq benchmarking dataset. We also found that higher biological diversity in pretraining data accelerated model enhancement and raised its performance limit in cell clustering.

Although stFormer was pretrained on spot-resolution Visium data, it is generalizable to ST data with single-cell resolution, such as CosMx SMI data. Our results indicated that spatial ligand expression information aided in cell type prediction for single-cell ST data. We provided an example to illustrate how cell type prediction benefited from a stable spatial niche.

In the gene-level tasks, we showed that our spatially aware gene embeddings more accurately predicted gene functions. It also uncovered intercellular ligand-receptor signaling responses through *in silico* perturbation of gene expressions, exemplified by the WNT signaling pathway in cardiac remodeling.

In summary, stFormer offers a flexible approach to spatial transcriptomics analysis. It learned general knowledge from pretraining ST data and transferred it to specific ST data for biological exploration. Although the current pretraining dataset was built on Visium data and remained relatively limited in scale, our model architecture was designed to be compatible with other types of ST data. As the volume of ST data in public databases continues to grow—particularly with the advancement of ST technologies that offer both single-cell resolution and high gene coverage—a larger-scale pretrained model can be developed in the future.

## Methods

### Model architecture

#### Assumptions

Suppose that the ST data we are analyzing are resolved at the single-cell level. Otherwise, if the ST data are generated through spot RNA sequencing, such as with the 10x Genomics Visium platform, we can follow the methods described in the next subsection to convert them to single-cell scenarios. Centered on each cell in ST data, the neighborhood within a preset radius is defined as the cell’s niche. For Visium platform, the niche radius is set to the spot radius. For single-cell ST data, such as CosMx SMI data, stFormer offers an option—*max number of niche cells* (*m*)—to define the niche radius as the maximum radius ensuring all cells have no more than *m* niche cells.

#### Layers

Schematic overview of the model architecture is illustrated in Fig. 1a. It is composed of six identical transformer blocks, each composed of a self-attention layer, cross-attention layer, and feed-forward neural network layer with the following parameters: 768 embedding dimensions, 12 attention heads per layer, and feed-forward size of 3072. Each layer includes attention computation or feed-forward neural network, residue connection, and layer normalization. stFormer can operate in single-cell mode, skipping cross-attention computation while retaining layer normalization in the cross-attention layer.

#### Input

The input sample consists of the gene symbols and values of the center cell, as well as the ligand gene symbols and values of the niche cells. We constructed a ligand gene library by integrating seven knowledge databases^17-23^ and only used the ligand genes included in at least two databases. We limit the input to genes with non-zero expressions. For each gene, the symbol embedding is retrieved from a learnable lookup table with a vocabulary size of 19,264. The embeddings in the lookup table are initialized by those from the scFoundation^5^ model. The gene value is transformed into a learnable value embedding through a scalar encoding module borrowed from scFoundation^5^. The final input embedding for each gene is the sum of its symbol embedding and value embedding.

#### Output

The transformer block first calculates self-attention among all gene embeddings within center cell, then computes cross-attention between all gene embeddings of center cell and ligand gene embeddings of niche cells (this step is skipped in single-cell mode), and finally propagates the gene embeddings through a two-layer feed-forward neural network. The model ultimately outputs gene embeddings specific to both the center cell’s intracellular context and spatial niche.

### Cell-type-wise biased cross-attention method on Visium data

Visium data consists of spot-wise gene expression profiles, with each spot covering multiple cells. We initially used the deconvolution method cell2location^7^ with default settings to disentangle the mixed RNA measurements at each spot. Cell2location outputs the proportions of different cell types and cell-type specific expression profiles at each spot. To utilize the model architecture described in the previous subsection, we developed the cell-type-wise biased cross-attention method based on the following two assumptions.

#### Assumption 1

Cells of the same type within a spot approximately have the same expression profiles. *Assumption 2*. The niche of each cell in a spot approximates to the cellular composition of that spot. The assumptions are reasonable since the spot size 55μm is relatively small, several-fold larger than a typical cell size, and *Assumption 1* is also adopted by most cell-type deconvolution methods. Under these two assumptions, calculating cross-attention using cell-wise ligand gene embeddings is equivalent to using cell-type-wise ligand gene embeddings with biases of logarithmic cell-type proportions, so that our model can do learning with single-cell resolution on the cell-type resolution Visium data. The details are described as follows.

Suppose that a total of *n* cells from *t* different cell types are located in a given spot, and the cell-type proportions estimated by cell2location are *p*_1_, *p*_2_, …, *p*_*t*_ respectively. For the sake of simplicity and without loss of generality, we suppose only one ligand gene *l* is detected. According to *Assumption 1*, the *np*_*i*_ cells of the *i*th cell type approximately have identical ligand gene expression, which is estimated by cell2location, resulting in the same ligand gene embedding, denoted as 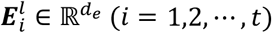.

For any one cell **c** in the spot, according to *Assumption 2*, the niche cells centered on cell **c** are approximately the *n* cells in the spot. Denote an arbitrary gene embedding of cell **c** as 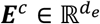. Then, the cross-attention between ***E***^*c*^ and the ligand gene embeddings of its niche cells 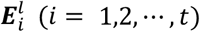 can be calculated as follows.

First, the gene embeddings are projected to the query, key, and value spaces respectively using the learnable parameter matrices 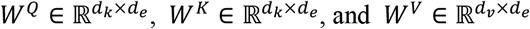.

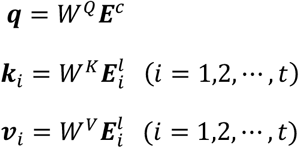

Then, the keys and values are packed together into matrices *K* and *V*, where 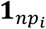 is an *np*_*i*_-dimensional vector with all elements being 1 and the prime symbol denotes matrix transpose.

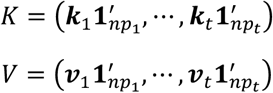

Finally, the attention function between ***q*** and *K, V* are computed as follows.

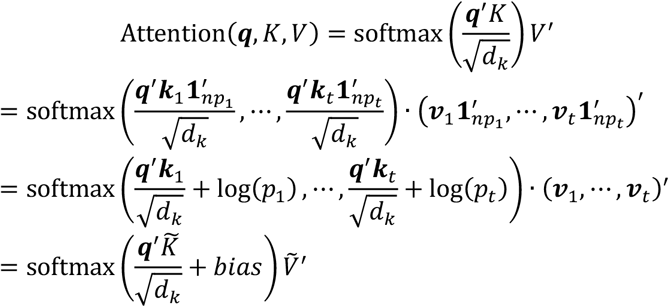

where 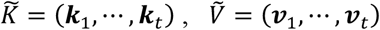 are the cell-type-wise keys and values; *bias* = (log(*p*_1_), …, log(*p*_2_)) is a bias term composed of logarithmic cell-type proportions. Detailed derivation of the above equations can be found in **Supplementary Note S1**.

### Pretraining

The pretraining was accomplished using a masked learning objective. Specifically, 15% of the genes in the center cells were masked. Their value embeddings were replaced with mask embeddings and then added with the symbol embeddings to form the input embeddings. A three-layer perceptron block was added after the model architecture to predict the expression values of the masked genes according to their output gene embeddings. The pretraining objective was to minimize the mean squared error (MSE) loss between the predicted and true values of the masked genes.

We assembled a pretraining corpus by deconvoluting 432 tissue sections from 75 human Visium datasets in CROST^8^, a public ST data repository. The corpus encompassed totally ~4.1 million samples across diverse tissues, development stages, and disease states (**Supplementary Data 1**). The model was pretrained with the following hyperparameters: epochs, 2; batch size, 4; gradient accumulation step, 2; learning rate 1 × 10^56^; optimizer, Adam. Pretraining lasted ~10 days on four Nvidia RTX 3090 24GB GPUs.

### Downstream tasks

#### Clustering of cell embeddings

The cell embedding for each cell was generated by max-pooling all of its output gene embeddings. We evaluated the batch effect on cell embeddings using a public benchmarking scRNA-seq dataset^9,10^, including 8 patients and 3 controls. In this case, stFormer was run in single-cell mode, and batch correction was quantified by ASW_batch_ score, calculated as inverse of the average silhouette width for batch clustering^11^. For comparison, we also evaluated the performance of a single-cell transcriptomics foundation model, scFoundation^5^.

We evaluated clustering results varying with pretraining data sizes on a normal myocardium section (ACH003) from a myocardial infarction Visium dataset^12^. To quantify the consistency between clustering results and annotated cell types, two metrics NMI (normalized mutual information) and ARI (adjusted Rand index) were calculated using the Python implementation of scib.metrics^11^. We constructed a less diverse pretraining corpus by deconvoluting a subset of Visium datasets (55 out of 75) and yielding ~2.4 million spatial samples. We also evaluated clustering results varying with pretraining data sizes on this reduced corpus. We evaluated clustering results before and after fine-tuning on a normal myocardium section (10×001) ^12^, and compared with scFoundation. Fine-tuning was conducted using the same objective function and hyperparameters as in pretraining solely on the test data.

### Cell type prediction

We attached a three-layer perceptron classifier to stFormer to perform the predictive task based on cell embeddings. The objective was to minimize the cross-entropy loss between predicted cell type probabilities and ground truth labels. We evaluated on a single-cell resolution CosMx SMI dataset from human pancreas tissue^14^. To avoid class imbalance effects, we downsampled all cell types to equal sample numbers. Then, the dataset was split into training (60%), validation (20%), and test (20%) sets. We fine-tuned the last two transformer layers together with the classifier. We compared two stFormer versions, stFormer-4.1M and stFormer-2.4M, at different niche radius settings, where *max number of niche cells* = 5, 10, 15, 20, 25, 30, and 35 respectively. stFormer-4.1M was pretrained on ~4.1 million samples while stFormer-2.4M was pretrained on ~2.4 million samples. For comparison, we also evaluated the predictive performance of scFoundation.

To visualize the spatial distribution of predictive results, we excluded one field-of-view (FOV=52) from the dataset, and downsampled remaining data to balanced class distribution. Then, we split training (80%) and validation (20%) sets, and performed model training and selection on them respectively. Finally, we predicted cell types on the held-out FOV, and compared the spatial distributions of acinar.1 cell predictions from stFormer and scFoundation.

### Gene function prediction

We attached a three-layer perceptron classifier to stFormer to predict gene inclusion within gene set based on gene embeddings. The objective was to minimize the cross-entropy loss between predicted gene inclusion probabilities and ground truth labels. The last two transformer layers were fine-tuned together with the classifier. We evaluated the predictive performance using fivefold cross-validation, and the results were reported as median ROC and PR curves with 40% to 60% quantile intervals. We evaluated on the KEGG gene set “TGF-beta signaling pathway” and the hallmark gene set “TNFA signaling via NFKB” provided by MSigDB^15^ (https://www.gsea-msigdb.org/gsea/msigdb/human/genesets.jsp). External signaling genes were excluded from the gene set, and an equal number of negative control genes were randomly sampled from outside the gene set. The evaluation samples were obtained from a fibrotic myocardium section (ACH005) in the myocardial infarction Visium dataset^12^. For comparison, we also evaluated the predictive performance of scFoundation.

### *In silico* perturbation analysis of ligand-receptor signaling responses

We employed stFormer to reveal the signaling responses of ligand-receptor interaction on downstream genes via *in silico* perturbation. We took the WNT signaling pathway in cardiac remodeling as an example. Specifically, we reduced the expression values of WNT ligand and receptor genes to one-third of baseline levels in cardiomyocytes on an ischemic tissue section (ACH0011) in the myocardial infarction Visium dataset^12^. Since we restricted the input to genes with non-zero expressions, the downregulated ligand genes were WNT2B, WNT5B and the downregulated receptor genes were FZD4, FZD5, FZD6. We analyzed the effects on various genes, including direct targets of the WNT signaling pathway, housekeeping genes (**Supplementary Table S1**), and 100 randomly selected genes, by calculating the cosine similarity of their output gene embeddings before and after *in silico* perturbation. Statistical comparisons of differential effects between gene groups were performed using the Wilcoxon rank-sum test, and the reported p-values were adjusted for multiple testing via false discovery rate (FDR) correction. The *in silico* perturbation was performed on ligand genes and receptor genes either individually or simultaneously.

## Data availability

The data used to produce the results were deposited on the Zenodo data repository under record number 16731623. The pretraining dataset is available upon request from the authors.

## Code availability

Models and weights, as well as the codes for pretraining and producing results are available on the Github repository at https://github.com/csh3/stFormer.

## Acknowledgments

This work was supported by the National Key R&D Program of China (2023YFF1204500), National Natural Science Foundation of China (No. 62103262).

## Author contributions

Shenghao Cao (Conceptualization, Investigation, Data curation, Formal analysis, Methodology, Visualization, Writing – original draft), Kaiyuan Yang (Methodology, Writing – review & editing), Jiabei Cheng (Methodology, Writing – review & editing), Jiachen Li (Methodology, Writing – review & editing), Hong-bin Shen (Supervision), Xiaoyong Pan (Supervision), Ye Yuan (Conceptualization, Methodology, Supervision, Project administration, Writing – review & editing)

