## Supplementary Information for "stFormer: a foundation model for spatial transcriptomics"

### Supplementary Notes

#### Note S1. Detailed derivation of the function $\mathrm{Attention}(\boldsymbol{q}, K, V)$ in the subsection "Cell-type-wise biased cross-attention method on Visium data" of the Methods section

$$K = \left(\boldsymbol{k}_1 \mathbf{1}'_{np_1}, \cdots, \boldsymbol{k}_t \mathbf{1}'_{np_t}\right) = (\underbrace{\boldsymbol{k}_1, \boldsymbol{k}_1, \cdots, \boldsymbol{k}_1}_{np_1}, \cdots, \underbrace{\boldsymbol{k}_t, \boldsymbol{k}_t, \cdots, \boldsymbol{k}_t}_{np_t})$$

$$V = \left(\boldsymbol{v}_1 \mathbf{1}'_{np_1}, \cdots, \boldsymbol{v}_t \mathbf{1}'_{np_t}\right) = (\underbrace{\boldsymbol{v}_1, \boldsymbol{v}_1, \cdots, \boldsymbol{v}_1}_{np_1}, \cdots, \underbrace{\boldsymbol{v}_t, \boldsymbol{v}_t, \cdots, \boldsymbol{v}_t}_{np_t})$$

$$\mathrm{Attention}(\boldsymbol{q}, K, V)$$

$$= \mathrm{softmax}\left(\frac{\boldsymbol{q}' K}{\sqrt{d_k}}\right) V'$$

$$= \mathrm{softmax}\left(\frac{\boldsymbol{q}' \boldsymbol{k}_1 \mathbf{1}'_{np_1}}{\sqrt{d_k}}, \cdots, \frac{\boldsymbol{q}' \boldsymbol{k}_t \mathbf{1}'_{np_t}}{\sqrt{d_k}}\right) \cdot \left(\boldsymbol{v}_1 \mathbf{1}'_{np_1}, \cdots, \boldsymbol{v}_t \mathbf{1}'_{np_t}\right)'$$

$$= \mathrm{softmax}\left(\frac{\boldsymbol{q}'(\boldsymbol{k}_1, \boldsymbol{k}_1, \cdots, \boldsymbol{k}_1)}{\sqrt{d_k}}, \cdots, \frac{\boldsymbol{q}'(\boldsymbol{k}_t, \boldsymbol{k}_t, \cdots, \boldsymbol{k}_t)}{\sqrt{d_k}}\right) \cdot (\underbrace{\boldsymbol{v}_1, \boldsymbol{v}_1, \cdots, \boldsymbol{v}_1}_{np_1}, \cdots, \underbrace{\boldsymbol{v}_t, \boldsymbol{v}_t, \cdots, \boldsymbol{v}_t}_{np_t})'$$

$$= \frac{\left(\exp\left(\frac{\boldsymbol{q}'\boldsymbol{k}_1}{\sqrt{d_k}}\right), \exp\left(\frac{\boldsymbol{q}'\boldsymbol{k}_1}{\sqrt{d_k}}\right), \cdots, \exp\left(\frac{\boldsymbol{q}'\boldsymbol{k}_1}{\sqrt{d_k}}\right), \cdots, \exp\left(\frac{\boldsymbol{q}'\boldsymbol{k}_t}{\sqrt{d_k}}\right), \exp\left(\frac{\boldsymbol{q}'\boldsymbol{k}_t}{\sqrt{d_k}}\right), \cdots, \exp\left(\frac{\boldsymbol{q}'\boldsymbol{k}_t}{\sqrt{d_k}}\right)\right)}{np_1 \times \exp\left(\frac{\boldsymbol{q}'\boldsymbol{k}_1}{\sqrt{d_k}}\right) + \cdots + np_t \times \exp\left(\frac{\boldsymbol{q}'\boldsymbol{k}_t}{\sqrt{d_k}}\right)} \cdot (\boldsymbol{v}_1, \boldsymbol{v}_1, \cdots, \boldsymbol{v}_1, \cdots, \boldsymbol{v}_t, \boldsymbol{v}_t, \cdots, \boldsymbol{v}_t)'$$

$$= \frac{np_1 \times \exp\left(\frac{\boldsymbol{q}'\boldsymbol{k}_1}{\sqrt{d_k}}\right) \cdot \boldsymbol{v}_1' + \cdots + np_t \times \exp\left(\frac{\boldsymbol{q}'\boldsymbol{k}_t}{\sqrt{d_k}}\right) \cdot \boldsymbol{v}_t'}{np_1 \times \exp\left(\frac{\boldsymbol{q}'\boldsymbol{k}_1}{\sqrt{d_k}}\right) + \cdots + np_t \times \exp\left(\frac{\boldsymbol{q}'\boldsymbol{k}_t}{\sqrt{d_k}}\right)}$$

$$= \frac{\exp\left(\frac{\boldsymbol{q}'\boldsymbol{k}_1}{\sqrt{d_k}} + \log(p_1)\right) \cdot \boldsymbol{v}_1' + \cdots + \exp\left(\frac{\boldsymbol{q}'\boldsymbol{k}_t}{\sqrt{d_k}} + \log(p_t)\right) \cdot \boldsymbol{v}_t'}{\exp\left(\frac{\boldsymbol{q}'\boldsymbol{k}_1}{\sqrt{d_k}} + \log(p_1)\right) + \cdots + \exp\left(\frac{\boldsymbol{q}'\boldsymbol{k}_t}{\sqrt{d_k}} + \log(p_t)\right)}$$

$$= \mathrm{softmax}\left(\frac{\boldsymbol{q}'\boldsymbol{k}_1}{\sqrt{d_k}} + \log(p_1), \cdots, \frac{\boldsymbol{q}'\boldsymbol{k}_t}{\sqrt{d_k}} + \log(p_t)\right) \cdot (\boldsymbol{v}_1, \cdots, \boldsymbol{v}_t)'$$

$$= \text{softmax}\left(\frac{\boldsymbol{q}'(\boldsymbol{k}_1, \cdots, \boldsymbol{k}_t)}{\sqrt{d_k}} + \big(\log(p_1), \cdots, \log(p_t)\big)\right) \cdot (\boldsymbol{v}_1, \cdots, \boldsymbol{v}_t)'$$

$$= \text{softmax}\left(\frac{\boldsymbol{q}'\widetilde{K}}{\sqrt{d_k}} + bias\right)\tilde{V}'$$

where $\widetilde{K} = (\boldsymbol{k}_1, \cdots, \boldsymbol{k}_t)$, $\tilde{V} = (\boldsymbol{v}_1, \cdots, \boldsymbol{v}_t)$, $bias = (\log(p_1), \cdots, \log(p_t))$.

### Supplementary Tables

**Table S1. WNT direct target genes and housekeeping genes analyzed to evaluate WNT signaling responses.**

| Symbol | Gene title | Class |
|---|---|---|
| FST | Follistatin | WNT direct target |
| MYC | Myc proto-oncogene protein | WNT direct target |
| MMP7 | Matrilysin | WNT direct target |
| MYCBP | c-Myc-binding protein | WNT direct target |
| FGF18 | Fibroblast growth factor 18 | WNT direct target |
| TCF4 | Transcription factor 4 | WNT direct target |
| NRCAM | Neuronal cell adhesion molecule | WNT direct target |
| JUN | Transcription factor Jun | WNT direct target |
| CCND1 | G1/S-specific cyclin-D1 | WNT direct target |
| ID2 | DNA-binding protein inhibitor ID-2 | WNT direct target |
| CLDN1 | Claudin-1 | WNT direct target |
| LEF1 | Lymphoid enhancer-binding factor 1 | WNT direct target |
| TBX3 | T-box transcription factor TBX3 | WNT direct target |
| AXIN2 | Axin-2 | WNT direct target |
| PPARD | Peroxisome proliferator-activated receptor delta | WNT direct target |
| VEGFA | Vascular endothelial growth factor A, long form | WNT direct target |
| FZD7 | Frizzled-7 | WNT direct target |
| FOSL1 | Fos-related antigen 1 | WNT direct target |
| TCF7 | Transcription factor 7 | WNT direct target |
| GAPDH | Glyceraldehyde-3-phosphate dehydrogenase | housekeeping |
| CHMP2A | Charged multivesicular body protein 2a | housekeeping |
| EMC7 | Endoplasmic reticulum membrane protein complex subunit 7 | housekeeping |
| GPI | Glucose-6-phosphate isomerase | housekeeping |
| PSMB2 | Proteasome subunit beta type-2 | housekeeping |
| PSMB4 | Proteasome subunit beta type-4 | housekeeping |
| RAB7A | Ras-related protein Rab-7a | housekeeping |
| REEP5 | Receptor expression-enhancing protein 5 | housekeeping |
| SNRPD3 | Small nuclear ribonucleoprotein Sm D3 | housekeeping |
| VCP | Transitional Endoplasmic Reticulum ATPase | housekeeping |
| VPS29 | Vacuolar protein sorting-associated protein 29 | housekeeping |

### Supplementary Figures

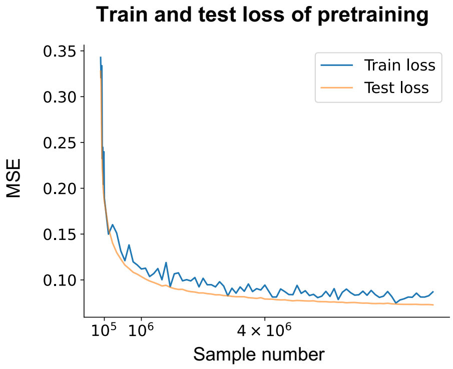

**Figure S1. Train and test loss during pretraining as the sample number increases, shown without log scaling.** MSE, mean squared error.

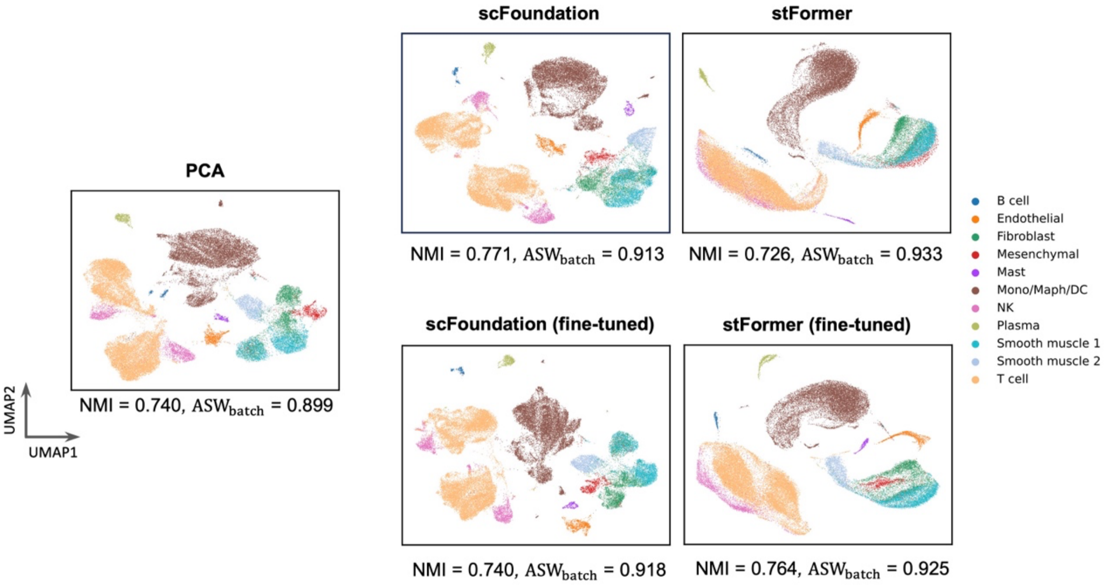

**Figure S2. Comparison for batch correction performance.** UMAP visualizations for cell embeddings generated on an scRNA-seq benchmarking dataset[1,2] by different approaches, with points colored by cell types. Clustering evaluation metric NMI and batch correction metric $ASW_{batch}$[33] are indicated below each UMAP plot. stFormer achieved the best performance for batch correction. After fine-tuning, stFormer showed a modest decline in $ASW_{batch}$, while its NMI markedly improved.

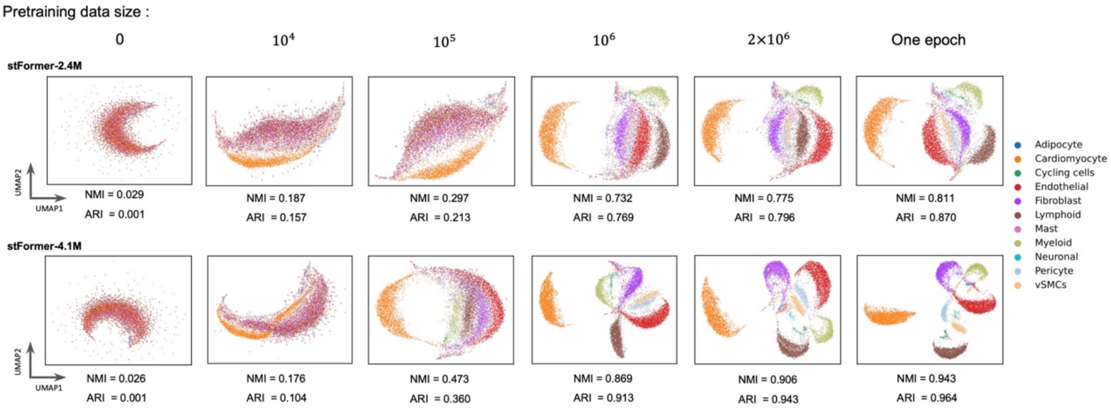

**Figure S3. UMAP visualizations for cell embeddings generated by stFormer-2.4 and stFormer-4.1 on a test Visium data as pretraining sample number increases.** Two clustering evaluation metrics, NMI (normalized mutual information) and ARI (adjusted Rand index), are indicated below each UMAP plot.

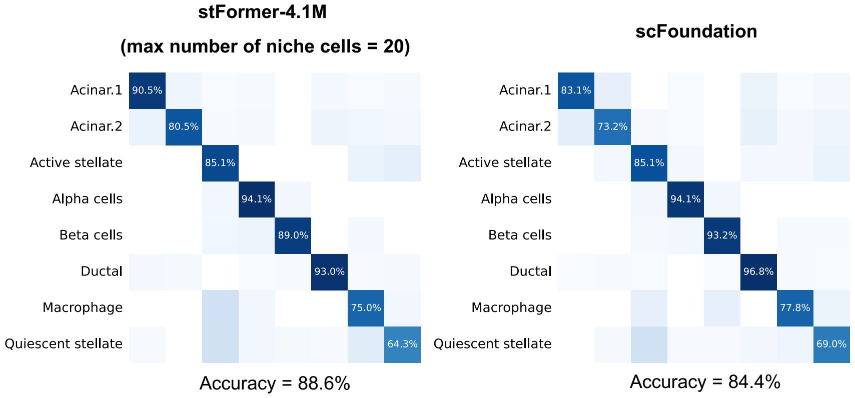

**Figure S4. Confusion matrices for the predictive results from stFormer-4.1M (*max number of niche cells* = 20) and scFoundation on the held-out FOV (FOV=52) of the human pancreas CosMx SMI dataset[4].** The overall predictive accuracy is indicated below each confusion matrix.

4 nanoString.
